# The Basal and Major Pilins in the *Corynebacterium diphtheriae* SpaA Pilus Adopt Similar Structures that Competitively React with the Pilin Polymerase

**DOI:** 10.1101/2023.02.23.529612

**Authors:** Christopher K. Sue, Nicole A. Cheung, Brendan J. Mahoney, Scott A. McConnell, Jack M. Scully, Janine Y. Fu, Chungyu Chang, Hung Ton-That, Joseph A. Loo, Robert T. Clubb

**Affiliations:** Department of Chemistry and Biochemistry, University of California, Los Angeles, 611 Charles Young Drive East, Los Angeles, CA 90095; UCLA-DOE Institute for Genomics and Proteomics, University of California, Los Angeles, 611 Charles Young Drive East, Los Angeles, CA 90095; Molecular Biology Institute, University of California, Los Angeles, 611 Charles Young Drive East, Los Angeles, CA 90095; Division of Oral and Systemic Health Sciences, School of Dentistry, University of California, Los Angeles, 611 Charles Young Drive East, Los Angeles, CA 90095

## Abstract

Many species of pathogenic gram-positive bacteria display covalently crosslinked protein polymers (called pili or fimbriae) that mediate microbial adhesion to host tissues. These structures are assembled by pilus-specific sortase enzymes that join the pilin components together via lysine-isopeptide bonds. The archetypal SpaA pilus from *Corynebacterium diphtheriae* is built by the ^Cd^SrtA pilus-specific sortase, which crosslinks lysine residues within the SpaA and SpaB pilins to build the shaft and base of the pilus, respectively. Here, we show that ^Cd^SrtA crosslinks SpaB to SpaA via a K139(SpaB)-T494(SpaA) lysine-isopeptide bond. Despite sharing only limited sequence homology, an NMR structure of SpaB reveals striking similarities with the N-terminal domain of SpaA (^N^SpaA) that is also crosslinked by ^Cd^SrtA. In particular, both pilins contain similarly positioned reactive lysine residues and adjacent disordered AB loops that are predicted to be involved in the recently proposed “latch” mechanism of isopeptide bond formation. Competition experiments using an inactive SpaB variant and additional NMR studies suggest that SpaB terminates SpaA polymerization by outcompeting ^N^SpaA for access to a shared thioester enzyme-substrate reaction intermediate.

## INTRODUCTION

Many bacterial species display pili (fimbriae), proteinaceous polymeric filaments that extend from the microbial surface to mediate cell-to-cell interactions, motility, and DNA uptake. Pathogens employ these structures during infections to adhere to host tissues, invade host cells, form biofilms, and modulate host immunity^1-5^. Many Gram-positive bacteria construct pili through a unique mechanism in which pilus-specific sortase enzymes connect the protein components of the pilus (called pilins) together via covalent lysine-isopeptide bonds that confer enormous tensile strength. These pili are composed of a single chain of crosslinked pilins that form long, thin hairlike structures which protrude 70-3000 nm from the cell surface^3^. A range of pathogenic bacterial species display sortase-crosslinked pili, including among others: *Streptococcus pyogenes, Streptococcus pneumoniae, Streptococcus agalactiae, Enterococcus faecalis, Clostridium difficile*, and *Corynebacterium diphtheriae*^6^. Commensal microbes that populate the human gut microbiome also display these structures to mediate microbe-microbe and microbe-host interactions, including: *Eubacterium lentum, Enterococcus faecium, Clostridium perfringens, Clostridium scindens*, and *Bifidobacterium infantis*^*7*^. As the rise of antibiotic resistant bacteria is an urgent and pressing medical problem, and pili are frequently virulence factors, a better understanding of how they are constructed could facilitate the development of pilus assembly inhibitors that function as antibiotics.

The mechanism of assembly of the *C. diphtheriae* SpaA pilus is paradigmatic^2,8-13^. It is constructed from three types of pilin subunits: SpaA, the major pilin that forms the pilus shaft; SpaB, a minor pilin that is located at the base of the pilus where it is attached to the cell wall, and SpaC, a minor pilin that is located at the pilus tip and mediates adherence to pharyngeal epithelial cells. While all pilins harbor a cell wall sorting signal (CWSS), SpaA contains a pilin motif with an active lysine residue required for crosslinking individual SpaA pilins^12^.In addition, SpaB possesses a critical lysine residue required for crosslinking the SpaA polymers to the basal SpaB pilin ^14^. These pilin components are assembled by two types of sortases, ^Cd^SrtA and ^Cd^SrtF. The ^Cd^SrtA enzyme is a pilus-specific sortase that covalently joins the SpaA, SpaB, and SpaC pilins together via lysine-isopeptide bonds, while ^Cd^SrtF attaches SpaB located at the base of the assembled pilus to the cell wall via a conventional peptide bond^14^. In the ^Cd^SrtA-catalyzed crosslinking reaction, the enzyme recognizes pilin substrates embedded in the membrane via a CWSS consisting of a LPXTG-motif followed by non-polar amino acids that form a transmembrane helix, and positively charged amino acids that are located at the C-terminal end of the protein and positioned within the cytoplasm. ^Cd^SrtA crosslinks the pilins by connecting the lysine residue in one pilin substrate to the threonine residue located within the LPXTG-motif of a second substrate. ^Cd^SrtA elongates the growing pilus by creating SpaA-x-SpaA crosslinks, where “-x-” represents the lysine-isopeptide bond, and the first and second pilins listed contribute reactive lysine and threonine residues, respectively (**Fig. 1A**). To install the SpaA-x-SpaA crosslink, the enzyme recognizes the lysine ε-amine group in one SpaA pilin (K190) and joins it the threonine carbonyl carbon atom in the LPLTG-motif located in another SpaA pilin (T494)^9^. A similar reaction is used to add the tip pilin via a SpaA-x-SpaC crosslink, but the threonine residue in this isopeptide bond originates from SpaC’s LPLTG-motif instead of SpaA^12^. Pilus polymerization is terminated by a distinct crosslinking reaction in which ^Cd^SrtA recognizes a lysine residue located in SpaB, enabling it to catalyze the formation of a SpaB-x-SpaA crosslink (**Fig. 1B)**^**14**^.

**Figure 1.**
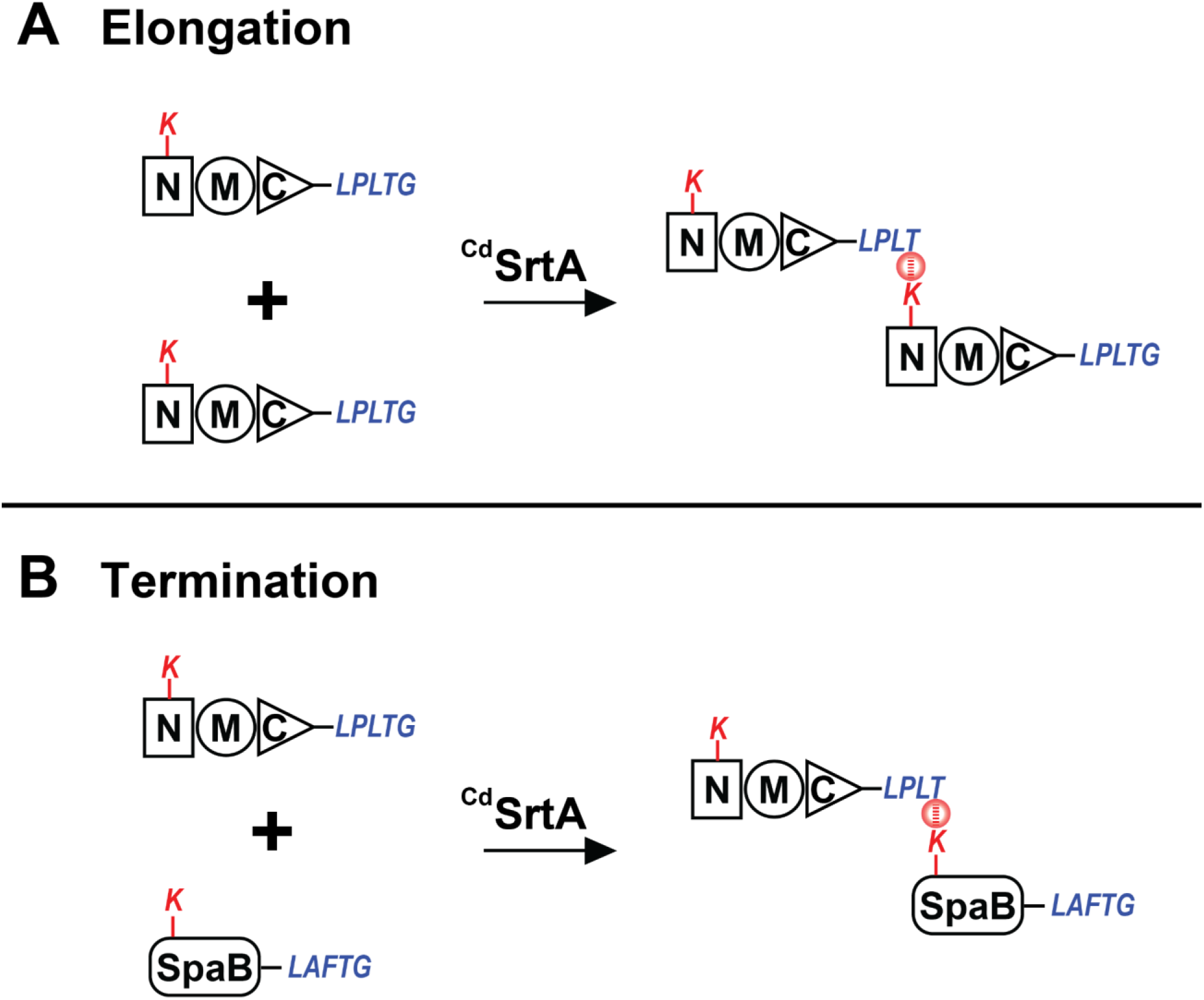
The lysine-isopeptide bond forming reactions catalyzed by ^Cd^SrtA. (A) Schematic showing the elongation reaction. Shown is the major SpaA pilin that contains N-terminal (N, square), middle (M, circle) and C-terminal (C, triangle) domains. SpaA proteins are joined when a lysine residue (K) within the N-terminal domain (red) is joined to the LPLTG sorting signal (blue). (B) Schematic showing the termination reaction. The LPLTG sorting signal within the major SpaA pilin (red) is joined to the lysine residue within the SpaB pilin (blue). In both images a circle containing a dashed line indicates the lysine-isopeptide bond.

Insight into the mechanism of SpaA-x-SpaA crosslinking has been gained from cellular, structural, and biochemical studies. A crystal structure of the intact SpaA pilin revealed that it contains three autonomously folded domains: N-terminal (^N^SpaA, residues 53 to 195), middle (^M^SpaA, residues 196 to 349), and C-terminal (^C^SpaA, residues 350 to 500) domains^10,13. Cd^SrtA installs interpilin SpaA-x-SpaA crosslinks by establishing a lysine–isopeptide bond between ^N^SpaA’s K190 ε-amine group and the T494 carbonyl group located within the LPLTG-motif that immediately follows ^C^SpaA in the primary sequence. SpaA-x-SpaA crosslinking has been reconstituted *in vitro*, enabling the determination of the NMR solution structure of the crosslinked ^N^SpaA-LPLT product complex in which K190 of ^N^SpaA is linked via an isopeptide bond to a peptide containing the LPLTG sorting signal^9,15^. Structures of the intact SpaA pilin and the product complex reveal that ^N^SpaA adopts a CnaB-type. The crosslinked sorting signal rests in a groove on ^N^SpaA that is formed by residues within a conserved WXXXVXVYPK pilin motif^12. Cd^SrtA has been proposed to use a latch mechanism to select K190 for crosslinking in a process that involves disordered-to-ordered transition in ^N^SpaA’s AB loop^10^. The structure of SpaB and the mechanism of its addition to the pilus to terminate its polymerization is not known. Here we show that despite sharing only 16% identity, SpaB and ^N^SpaA possess similar structures with features compatible with them being crosslinked via a recently proposed “latch mechanism”^10^. Kinetic measurements of crosslinking and NMR experiments suggest that the SpaB terminates pilus polymerization by selectively binding to only the thioester-linked ^Cd^SrtA-SpaA enzyme-substrate and not to the apo-form of ^Cd^SrtA.

## Materials and Methods

### Protein expression and purification

The protein expression plasmids were constructed using standard methods^15. C^SpaA constructs contained an N-terminal 6xHis tag in a pMAPLE4 expression vector. All other protein constructs were cloned into a pSUMO vector, which was modified from a pET-28b vector by insertion of an N-terminal small ubiquitin-like modifier (SUMO) solubility domain with a 6xHis tag. Proteins were expressed in *Escherichia coli* BL21 (DE3) cells. Cultures were grown in LB Miller broth (Thermo Fisher) supplemented with 50 µg/mL of kanamycin sulfate (GoldBio) at 37°C in a shaking incubator to the log growth phase (OD_600_ of 0.6 to 0.8) before being induced with 1 mM isopropyl β-d-1-thiogalactopyranoside (IPTG) (GoldBio). Cultures were then moved to low temperature orbital shakers and incubated at 17°C for 16 hours. For isotopically labeled samples, cell pellets were exchanged into M9 media supplemented with ^13^C-labeled glucose and ^15^N-labeled ammonium chloride before being induced with IPTG. Cells were then pelleted by spinning in a centrifuge at 8,500 x *g* for 15 min. Pellets were then redissolved in 50 mM Tris-HCl and 300 mM NaCl, at pH 8.0 (Buffer A). Subsequently, the cells were lysed using an Emulsiflex homogenizer and then fractionated via centrifugation at 22,720 *x* g for 50 min. Afterward, the cell lysate was purified via IMAC-Co^2+^ purification. Proteins were eluted from the resin using a lysis buffer supplemented with 200 mM imidazole. The 6xHis-SUMO tags were removed via treatment by 6xHis-Ulp1 protease at 1 mg/mL and subsequent purification by IMAC-Co^2+^. Afterward, the protein was purified by size exclusion chromatography using an AKTA Pure FPLC system (GE) and a column packed with Superdex 75 PG resin. Protein purity was confirmed by SDS-PAGE. pSUMO expression plasmids encoding ^Cd^SrtA^3M^ variants (residues N37–Q257 in ^Cd^SrtA with D81G/W83G/N85A substitutions in the lid) were created using standard molecular biology methods and confirmed by nucleotide sequencing^16^. All purified enzymes were stored at -20°C in Buffer A supplemented with 40% glycerol. The FELPLTGGSG peptide (LPLTG peptide) used in the transpeptidation and hydrolysis assays was synthesized by Peptide 2.0.

### Transpeptidation assays

Gel-based assays to measure crosslinking were performed as previously described^15,16^. In this assay, ^C^SpaA containing a CWSS was ligated to either SpaB or ^N^SpaA by ^Cd^SrtA^Δ^ and reactions were separated by SDS-PAGE. Reactions were performed in 100 μL volumes and contained 100 µM of ^Cd^SrtA^Δ^ (residues N37–Q257 of ^Cd^SrtA in which the amino acids I78 and A88 are deleted), 5 mM DTT, 300 µM of ^C^SpaA, and 300 µM of SpaB or ^N^SpaA. All components were dissolved in Buffer A, and reactions were initiated by adding the ^Cd^SrtA^Δ^ enzyme^9^. Reactions were conducted at 25°C and timepoints were taken at 0, 24, 48, and 72 hours. Samples were then separated by SDS-PAGE. An HPLC-based assay was used to quantify the kinetics of ^Cd^SrtA^Δ^ when SpaB is used as a substrate. In this assay, SpaB containing the reactive lysine is ligated to a FELPLTGGSG peptide (LPLTG peptide) by ^Cd^SrtA^Δ^ and reactions are HPLC-quantified using a C4 column (5 µm, 4.6 × 150 mm; Phenomenex). Reactions were performed in 100 μL volumes and contained 25 µM of ^Cd^SrtA^Δ^, 5 mM of DTT, 1 mM of LPLTG peptide, and variable amounts of SpaB (62.5 to 500 μM). All components were dissolved in Buffer A, and reactions were initiated by the addition of ^Cd^SrtA^Δ^, incubated for 3 h at 25°C, and then flash-frozen with liquid N_2_ and stored at -20°C. The reactions were analyzed using HPLC (Agilent 1100; C4 column: 5 μm, 4.6 × 150 mm; Phenomenex) with an initial dwell time of 3 min at 30% CH_3_CN/0.1% TFA followed by a linear gradient from 30% to 40% CH_3_CN/0.1% TFA for 23 min at 1 mL/min. The column was subsequently flushed with high concentrations of CH_3_CN/0.1% TFA at 1 mL/min. SpaB-containing peaks were detected at 215 nm and the amount of substrate converted to product was calculated by integrating the HPLC peak areas. The identity of each peak in the HPLC chromatogram was confirmed via LC-MS. Kinetic parameters of the reactions were obtained by fitting the data with SigmaPlot 12.0.

The pilin competition experiment was conducted with similar concentrations as above except reactions contained 200 µM of either ^N^SpaA, SpaB-APNV, and/or SpaB^K139^. Reaction samples were taken at 1, 3, 6, and 24 hours and quenched by freezing at -20°C prior to HPLC analysis. To analyze the samples, HPLC was used (Phenomenex C4 column: 5 μm, 4.6 × 150 mm) with a 3-minute dwell time of 30% CH_3_CN/0.1% TFA and subsequent linear gradient from 30% CH_3_CN/0.1% TFA to 44% CH_3_CN/0.1% TFA applied over 28 minutes with a flow rate of 1 mL/min. The column was then flushed with high concentrations of CH_3_CN/0.1% TFA to remove any remaining protein. Peaks corresponding to ^N^SpaA and SpaB were detected at 215 nm and quantified by integration. The amount of expected product was calculated assuming the reaction shown in Scheme 1 using the program MATLAB^17^.

### NMR experiments and structure determination

NMR spectra were collected at 298 K on Bruker Avance III HD 600 MHz and Avance NEO 800 MHz spectrometers equipped with triple resonance probes as previously described^10,18^. NMR data were processed with NMRPipe and analyzed using CARA (version 1.8.4), XIPP (version 1.19.6 p0), and NMRFAM-Sparky^19-21. 1^H, ^13^C, and ^15^N protein chemical shifts were assigned using the following experiments: ^15^N-HSQC, ^13^C-HSQC, HNCACB, CBCA(CO)NH, HNCO, HN(CA)CO, HBHA(CO)NH, HNHA, HNHB, CC(CO)NH, H(CCCO)NH, HCCH-COSY, HCCH-TOCSY, and ^15^N-TOCSY^22^. {^1^H}^15^N heteronuclear NOE experiments were acquired in duplicate and used to define regions of relative disorder, with data analyzed in NMRFAM-Sparky. ^15^N- and ^13^C-edited NOESY spectra (120 ms mixing time) were also acquired, and used to define NOE distance restraints^23^. The ϕ and ψ dihedral angle restraints were obtained using the program TALOS-N^24^. Structures were determined using the program XPLOR-NIH^25,26^. Initially, NOE cross peaks in the 3D ^15^N-edited NOESY-HSQC and ^13^C-edited NOESY-HSQC spectra were assigned automatically using the program UNIO10^27,28^. The NOESY data were then manually inspected using the program Xipp to verify all cross-peak assignments and to identify additional distance restraints^20^. An iterative procedure was used to refine the structure. In the final round of calculations, 200 structures were generated, which yielded a total of 50 with no NOE, or dihedral angle violations greater than 0.5 Å, 5°, or 2 Hz, respectively. The structures were sorted based on lowest overall energy, and the top 20 were selected as the ensemble to represent the structure of SpaB and have been deposited in the PDB. The programs MOLMOL and PyMOL were used to generate figures^29,30^.

### Tandem mass spectrometry

Protein digestion and isopeptide bond identification were performed according to a previously described protocol^15,16^. Specifically, proteins entrapped in gel bands were reduced with 10 mM dithiothreitol (Sigma) at 60°C for one hour and then alkylated with 50 mM iodoacetamide (Sigma) at 45°C for a few minutes in the dark. Samples were then digested with 200 ng trypsin (Thermo Scientific) at 37°C overnight. At the end of trypsin digestion, zinc acetate was added to the solution to a final concentration of 2.5 mM. 200 ng of Asp-N endoproteinase (Thermo Scientific) was then added for another overnight incubation. Digested peptides were extracted from the gel bands in 50% acetonitrile/49.9% water/0.1% trifluoroacetic acid (TFA) and cleaned with C18 StageTips before mass spectrometry analysis^31^. Digested peptides were separated on an EASY-Spray column (25 cm × 75 μm ID, PepMap RSLC C18, 2 μm, Thermo Scientific) connected to an EASY-nLC 1000 nUPLC (Thermo Scientific) using a gradient of 5-35% acetonitrile with 0.1% formic acid, and a flow rate of 300 nl/min (total time 30 minutes). Tandem mass spectra were acquired in a data-dependent manner with an Orbitrap Q Exactive mass spectrometer (Thermo Fisher Scientific) interfaced to a nanoelectrospray ionization source. The raw MS/MS data were converted into MGF format by Thermo Proteome Discoverer 1.4 (Thermo Scientific). An in-house program was used to search for the isopeptides. The approach was based on the observation of published spectra as well as our own on the presence of ions specific for the fragments of FELPLT (m/z 249.12341, 277.11833, 390.2023).

## Results

### The ^Cd^SrtA polymerase joins SpaB to SpaA via a K139-T494 lysine-isopeptide bond

*In vivo* studies have shown that K139 in SpaB is essential for its incorporation into the pilus and that this residue is important for ^Cd^SrtA-catalyzed crosslinking of SpaB to SpaA *in vitro*^32^. However, the specific residues that participate in the SpaB-x-SpaA crosslink produced by SrtA have not been identified experimentally. To investigate this issue, we reconstituted the SpaB-x-SpaA crosslinking reaction *in vitro* using ^Cd^SrtA^Δ^, an activated form of enzyme in which several residues within its inhibitory lid structure are removed^9^. As shown previously and in **Fig. 2A**, incubating ^Cd^SrtA^Δ^ with polypeptides containing SpaA’s N-terminal (^N^SpaA, residues 53 to 195) and C-terminal (^C^SpaA, residues 350-500 domains of SpaA containing a LPLTG sorting signal) domains results in the appearance of a crosslinked product (^N^SpaA-x-^C^SpaA). This reaction represents a single ligation event in which ^Cd^SrtA^Δ^ links adjacent SpaA pilins via a K190(^N^SpaA)-T494(^C^SpaA) lysine-isopeptide bond. As shown in **Figs. 2B** and **2C** (left panels), when ^Cd^SrtA^Δ^ is incubated with ^C^SpaA and a polypeptide encoding the SpaB protein lacking its N-terminal secretion signal and CWSS (SpaB, residues 25–180), a new band at ∼30 kDa appears. LC-MS/MS analysis of this band confirms that it is the SpaB-x-^C^SpaA crosslinked product formed by ^Cd^SrtA^Δ^ in which the ε-amine of K139 in SpaB is joined via an isopeptide bond to T494 within the LPLTG sorting signal that follows the ^C^SpaA domain (**Fig. 2D**). SpaB contains three lysines: K53, K115, and K139. ^Cd^SrtA^Δ^ is highly selective for the K139 side chain as only variants in which it is exchanged with alanine are refractory to catalysis (**Fig. 2C**, right), whereas K53A and K115A variants are crosslinked to SpaA similar to the wild-type protein (**Fig. 2B**). Thus, our results demonstrate that ^Cd^SrtA^Δ^ joins SpaB to the base of the pilus by selectively catalyzing the formation of the K139(SpaB)-T494(^C^SpaA) lysine-isopeptide bond.

**Figure 2.**
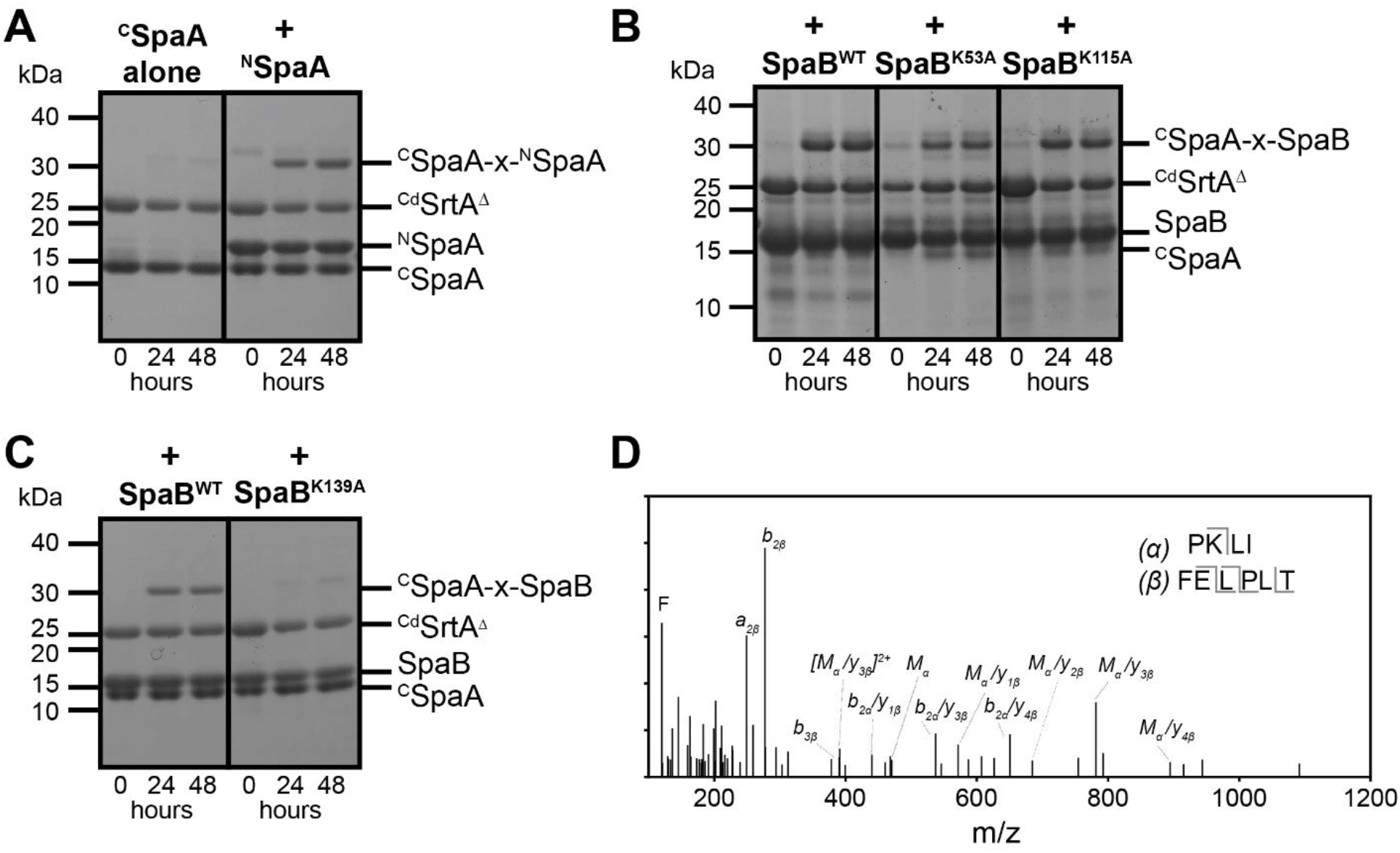
^Cd^SrtA selectively uses K139 within SpaB for crosslinking. (A) *In vitro* reconstitution of polymerase catalyzed ^N^SpaA-^C^SpaA crosslinking reaction. The reactions contained ^Cd^SrtA^Δ, N^SpaA, and ^C^SpaA at concentrations of 100 µM, 300 µM, and 300 µM, respectively. Reactions were incubated at 25°C for 0, 24, and 48 hours (left, middle, and right of each panel). (B) *In vitro* reconstitution of the SpaB-^C^SpaA crosslinking reaction. Reactions were performed as in panel (A), except that ^N^SpaA was replaced with wild-type SpaB (left, SpaB^WT^) or SpaB containing K53A (middle, +SpaB^K53A^) or K115A (right, +SpaB^K115A^) amino acid substitutions. (C) Reconstituted reactions containing ^Cd^SrtA^Δ^ and ^C^SpaA, and either wild-type SpaB (left, +SpaB^WT^) or the K139A variant (right, +SpaB^K139A^). (D) Mass spectrometry (LC-MS/MS) identification of the residues used to form the lysine isopeptide bond between SpaA and SpaB. The panel shows the fragmentation spectra of the linked peptide to SpaB K139 (sequence shown in insert, α is the peptide sequence from SpaB and β is the synthetic LPLTG peptide sequence). Detected fragment ions (M, a-, b-, and y-ions) are labeled accordingly and reported in Supplemental Table S1.

### Kinetic measurements of SpaB-x-SpaA crosslinking

We previously measured the *in vitro* kinetics of SpaA-x-SpaA crosslinking using an HPLC-based assay that monitors the rate at which ^Cd^SrtA joins the ^N^SpaA domain to a peptide derived from ^C^SpaA’s CWSS (FELPLTGGSG, hereafter referred to as “LPLTG”)^9^. Here, we extended this assay to quantify the kinetics of SpaB-x-SpaA crosslinking that terminates pilus assembly (**Fig. 3A**). As shown in Scheme 1, the first step in the SpaA-x-SpaA and SpaB-x-SpaA crosslinking reactions are identical and involve formation of a ^Cd^SrtA-LPLT thioester-linked enzyme-substrate intermediate (ES’).

**Scheme 1.**
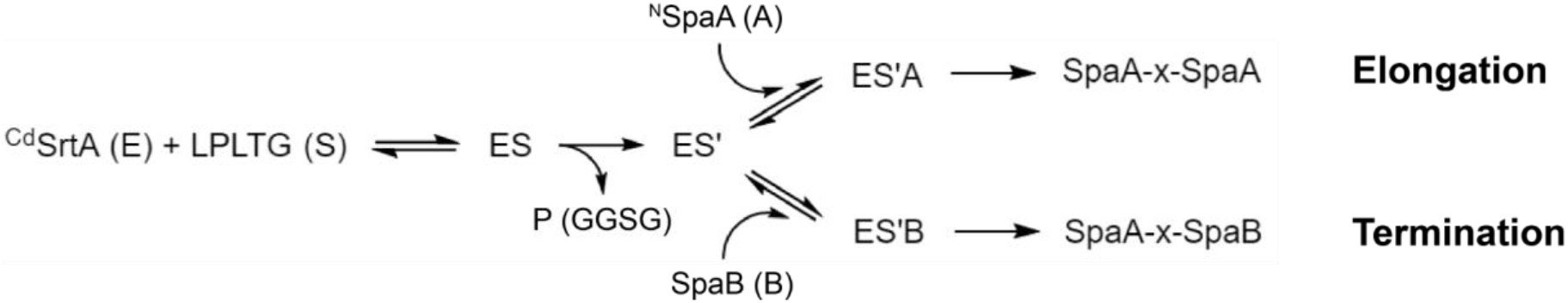

**Figure 3.**
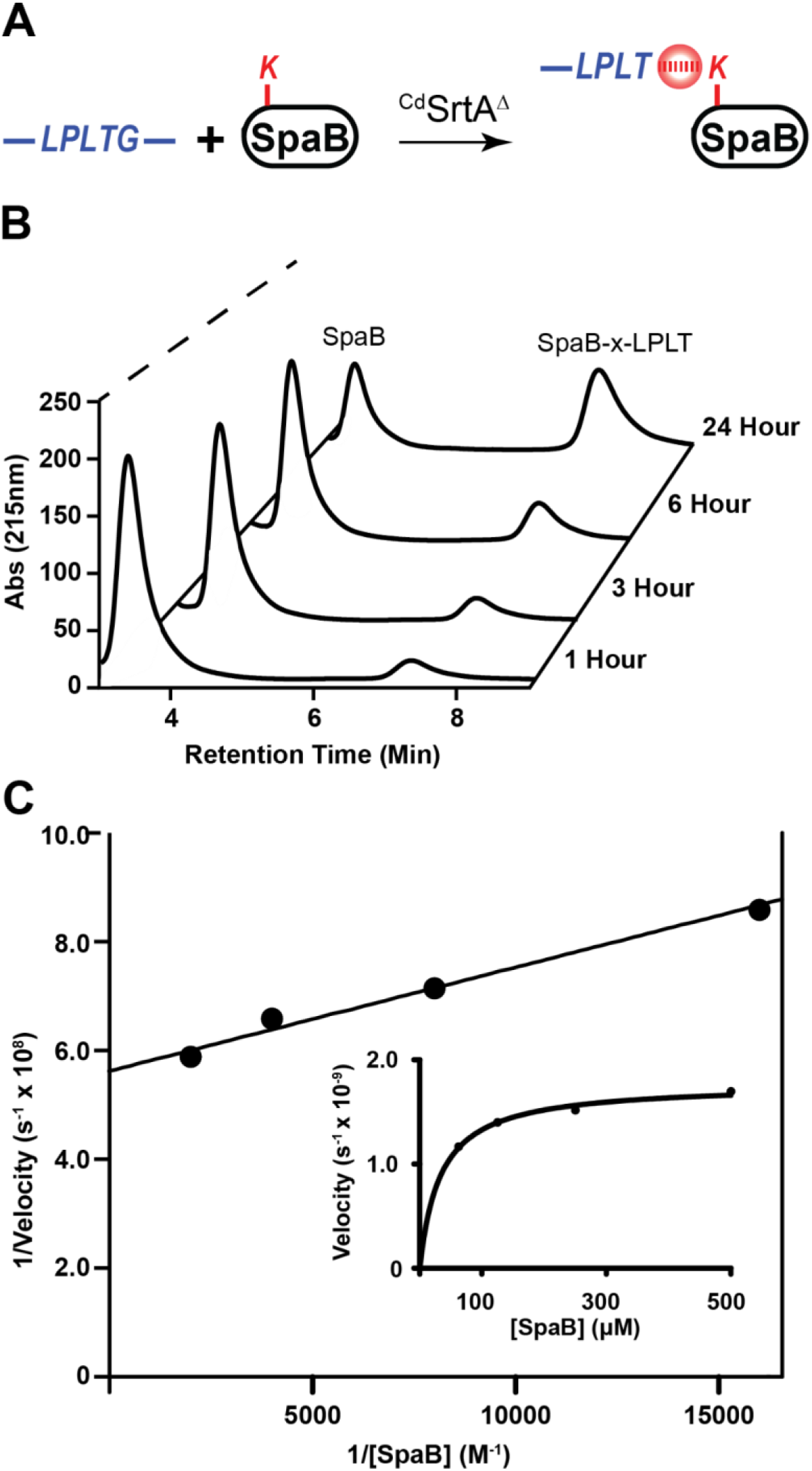
Kinetic measurements of SpaB crosslinking. (A) Schematic showing the termination reaction catalyzed by ^Cd^SrtA^Δ^ that joins the LPLTG peptide within the CWSS of SpaA to K139 within the SpaB basal pilin. (B) Overlaid HPLC traces showing reaction progress as a function of time. The reactions contained: 25 μM enzyme, 100 μM SpaB, and 1 mM LPLTG peptide. The amount of SpaB reactant and SpaB-x-LPLT product were quantified. (C) Linear fitting of the Lineweaver-Burk plot of reciprocal initial velocities versus reciprocal SpaB concentration. The data were acquired in triplicate. The insert shows the same data, but with the initial velocity versus substrate SpaB concentration plotted.

The thioester intermediate (ES’) forms when C222 within ^Cd^SrtA nucleophilically attacks the carbonyl group located in the threonine residue of the LPLTG sorting signal, which cleaves its threonine-glycine peptide bond leaving the threonine portion of the peptide attached to the enzyme. The ES’ intermediate is then poised to react with lysine residues located in either ^N^SpaA (K190) or SpaB (K139) to produce their respective crosslinked products. To probe the kinetics of the second step in the reaction that is unique to each crosslinking reaction we measured the rates of product formation when excess LPLTG peptide was present (1 mM). **Figure 3B** shows representative HPLC data revealing that the amount of SpaB reactant and isopeptide-linked SpaB-x-LPLT product can be quantified using reversed phase HPLC. Velocity measurements as a function of SpaB concentration were made as previously described (62.5 to 500 µM SpaB). An analysis of the steady state kinetics data yielded a K_M_ for SpaB (^SpaB^K_M_) of 3.2 ± 0.2 µM and a k_cat_ of 7.0 ± 0.4 × 10^−5^ s^-1^ (^SpaB^k_cat_) (**Fig. 3C**). We have previously measured the kinetics of ^Cd^SrtA^Δ^ catalyzed crosslinking of ^N^SpaA using identical reaction conditions, except that ^N^SpaA was used instead of SpaB. ^Cd^SrtA^Δ^ crosslinks ^N^SpaA to the LPLTG peptide with a measured K_M_ for the ^N^SpaA substrate (^SpaA^K_M_) of 16 ± 4 µM and a k_cat_ of 40 ± 0.1 × 10^−5^ s^-1^ (^SpaA^k_cat_). Thus, based on their K_M_ values, ^Cd^SrtA^Δ^ presumably has higher affinity for SpaB as compared to ^N^SpaA, but when the enzyme is saturated with its substrates, it crosslinks the sorting signal to SpaB ∼6-times slower than to ^N^SpaA.

### SpaB interacts with ^Cd^SrtA-LPLT thioester intermediate to slow SpaA-x-SpaA crosslinking

SpaA pili become elongated when either SpaA is overexpressed or the amount of SpaB is reduced^12,14^. This suggests that on the cell surface the SpaA and SpaB pilins stochastically compete with one another for access to ^Cd^SrtA to either continue or terminate pilus assembly, respectively. We investigated this issue by monitoring the *in vitro* effect of SpaB on the ^N^SpaA-LPLT crosslinking reaction. Initially, separate crosslinking experiments were performed in which ^Cd^SrtA^Δ^ and the LPLTG peptide were incubated with either ^N^SpaA or SpaB. In these experiments excess LPLTG peptide was present to ensure that product formation was primarily dependent on the second step of catalysis (nucleophilic attack of either ^N^SpaA or SpaB on the ^Cd^SrtA-LPLT thioester intermediate) (Scheme 1). **Figure 4A** shows the fraction of ^N^SpaA or SpaB protein converted to ^N^SpaA-x-LPLT and SpaB-x-LPLT product, respectively. As expected, the rate at which the enzyme catalyzes LPLTG peptide modification of ^N^SpaA is faster than for SpaB, with ∼58.3 +/- 0.6% and 24.3 +/- 0.2% of these proteins converted to product after 6 hours, respectively. Simulations of the modification reactions that assume saturating concentrations of the sorting signal and the mechanism outlined in Scheme 1 are in good agreement with the experimental data (dashed lines in **Fig. 4A**). Good agreement with the experimental data is obtained when the simulations employed K_M_ and k_cat_ values similar to those obtained from the Lineweaver-Burk analysis. Similar reactions were performed to determine the effects of SpaB on the rate of ^N^SpaA crosslinking by ^Cd^SrtA^Δ^. When the ^N^SpaA crosslinking is repeated in the presence of an equimolar amount of wild type SpaB a marked drop in ^N^SpaA-x-LPLT product formation occurs (**Fig. 4B**). This is compatible with both types of pilins reacting with the ^Cd^SrtA-LPLT thioester intermediate, which is depleted as a result of SpaB binding (Scheme 1). To quantify the affinity of SpaB for the intermediate, the ^N^SpaA crosslinking reaction was repeated in the presence of SpaB^K139A^, which is unreactive and presumably only binds to the enzyme (**Fig. 4C**). Interestingly, a large reduction in ^N^SpaA-x-LPLT product formation occurs in the presence of the SpaB mutant. Modeling the reaction assuming SpaB^K139A^ acts as a competitive inhibitor that binds to the ^Cd^SrtA-LPLT thioester intermediate yields a K_D_ ∼65 µM.

**Figure 4.**
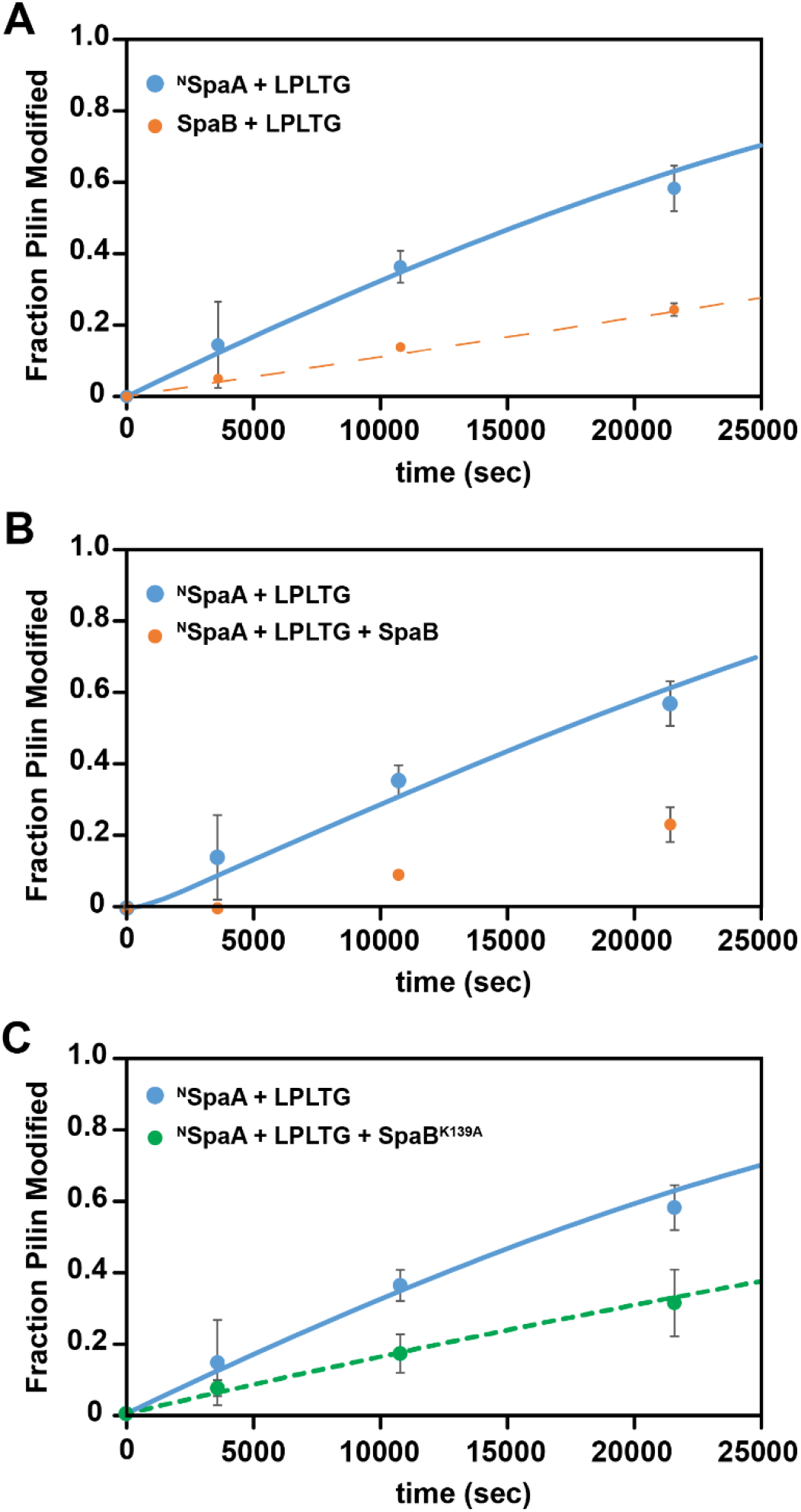
*In vitro* crosslinking competition between ^N^SpaA and SpaB. The panels show time courses of crosslinking reactions containing ^N^SpaA and wild-type and mutant SpaB proteins. (A) A plot of the time course of pilin protein crosslinking by ^Cd^SrtA^Δ^ with the LPLTG peptide. The fraction of ^N^SpaA (blue) and SpaB (orange) converted to their crosslinked product is shown. In the reactions 25 μM ^Cd^SrtA^Δ^ and 4 mM LPLTG peptide were incubated with either ^N^SpaA or SpaB (200 μM) and the fraction of pilin protein that was crosslinked with the peptide was determined by HPLC. (B) Same as panel (A) showing the fraction of ^N^SpaA converted to its crosslinked product in the absence (blue) or presence of 200 μM SpaB (orange). In the reactions the concentration of ^N^SpaA was 200 μM. (C) Same as panel (B), except that instead of wild type SpaB, an unreactive SpaB^K139A^ variant was employed. The mutant does not react with the enzyme, but nevertheless inhibits SpaA crosslinking. The error bars for the experimental data are the standard deviations of the measured values obtained from three experiments. The dashed lines are the predicted values obtained from simulations (described in the Methods section).

The results of the competition experiments and the enzyme’s smaller K_M_ value for SpaB suggest that SpaB slows the SpaA-x-SpaA crosslinking by outcompeting ^N^SpaA for access to the ^Cd^SrtA-LPLT thioester intermediate. However, SpaB could also slow SpaA-x-SpaA crosslinking by binding to either the apo-form of the enzyme or to the LPLTG peptide. This slower reaction caused by binding to the LPLTG peptide seems unlikely because the peptide is in significant excess. NMR was used to probe the strength of SpaB binding to the apo-form of the enzyme (similar studies using the ^Cd^SrtA-LPLT thioester intermediate are not possible because it is too sparsely populated in solution). The backbone chemical shifts of SpaB were assigned using triple-resonance NMR approaches. The ^1^H-^15^N HSQC spectrum of ^15^N-labeled SpaB is well resolved enabling the backbone chemical shifts of 89.7% of its residues to be assigned (**Fig. 5A**). A series of titration experiments were performed in which ^1^H-^15^N HSQC spectra of ^15^N-labeled SpaB were recorded in the presence of varying amounts of unlabeled ^Cd^SrtA^2M^, an enzymatically active form of ^Cd^SrtA that contains two destabilizing mutations in its inhibitory lid structure (residues P77-S89 of ^Cd^SrtA containing D81G and W83G substitutions). As shown in **Figs. 5B** and **5C**, adding the enzyme to SpaB does not alter its NMR spectrum, indicating that the proteins do not interact with one another with high affinity. Even at 10:1 enzyme:SpaB ratios only minimal spectral changes occur indicating that SpaB binds to the free enzyme with a K_D_ > ∼2 mM. Thus, we conclude that SpaB slows the SpaA-x-SpaA crosslinking reaction by preferentially binding to the ^Cd^SrtA-LPLT thioester intermediate.

**Figure 5.**
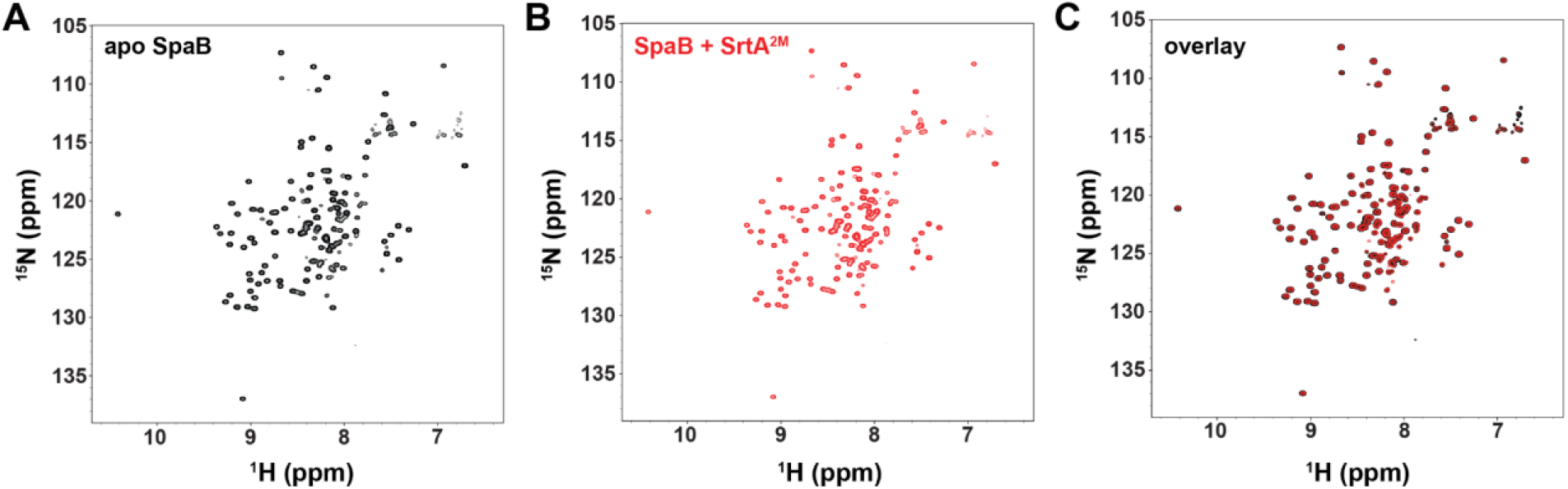
SpaB binds to the apo-form of the pilin-specific sortase with very weak affinity. ^1^H-^15^N HSQC spectra of SpaB (A) alone, (B) in the presence of unlabeled ^Cd^SrtA^2M^, and (C) an overlay of the spectra shown in panels (A) and (B). Spectra were recorded using 50 µM ^15^N-labeled SpaB and 350 µM unlabeled ^Cd^SrtA^2M^. No significant changes in SpaB’s spectrum occur upon adding the enzyme. This data indicates that the SpaB pilin does not interact with the apo-form of the enzyme with appreciable affinity (K_D_ > ∼2 mM).

### Structure of SpaB

To gain insight into how SpaB is used by ^Cd^SrtA as a substrate we determined its structure using NMR (**Table S2**). The backbone and heavy atom coordinates of residues N28-P37 and D51-D142 in the ensemble of 20 conformers are structurally ordered and can be superimposed to the average structure with a root mean square deviation (rmsd) of 0.47 ± 0.09 and 0.82 ± 0.07 Å, respectively (**Fig. 6A**). SpaB adopts a CnaB fold, an immunoglobulin-like domain first visualized in the staphylococcal collagen adhesin (Cna)^33-36^. It contains seven β-strands that form a β-sandwich, whose sides are composed of three- (strands G, A, and D) and four-stranded (strands C, B, E and F) β-sheets (**Fig. 6B**). The fold is supplemented by a single α-helix located in the polypeptide segment that connects strands B to C. The K139 side chain that is crosslinked by ^Cd^SrtA is positioned near the C-terminal end of the domain in an extended segment that follows strand G. The lysine side chain is positioned between the α-helix and a large structurally disordered loop that connects strands A to B (AB loop, residues 34-49). An analysis of the {^1^H}^15^N heteronuclear NOE (hetNOE) data for the backbone amide atoms within SpaB reveals that the loop is dynamic on the pico- to nano-second time scales as its residues have hetNOE values <0.6 (**Fig. 5C**). Residues E82, Q84, T113 are also flexible based on hetNOE data and are located within surface loops connecting α-helix to strand C (E82 and Q84), and strand E to F (T113). Notably, the region containing K139 (D128 through T135) has slightly elevated mobility, compatible with these residues located in an extended segment near the protein’s C-terminus.

**Figure 6.**
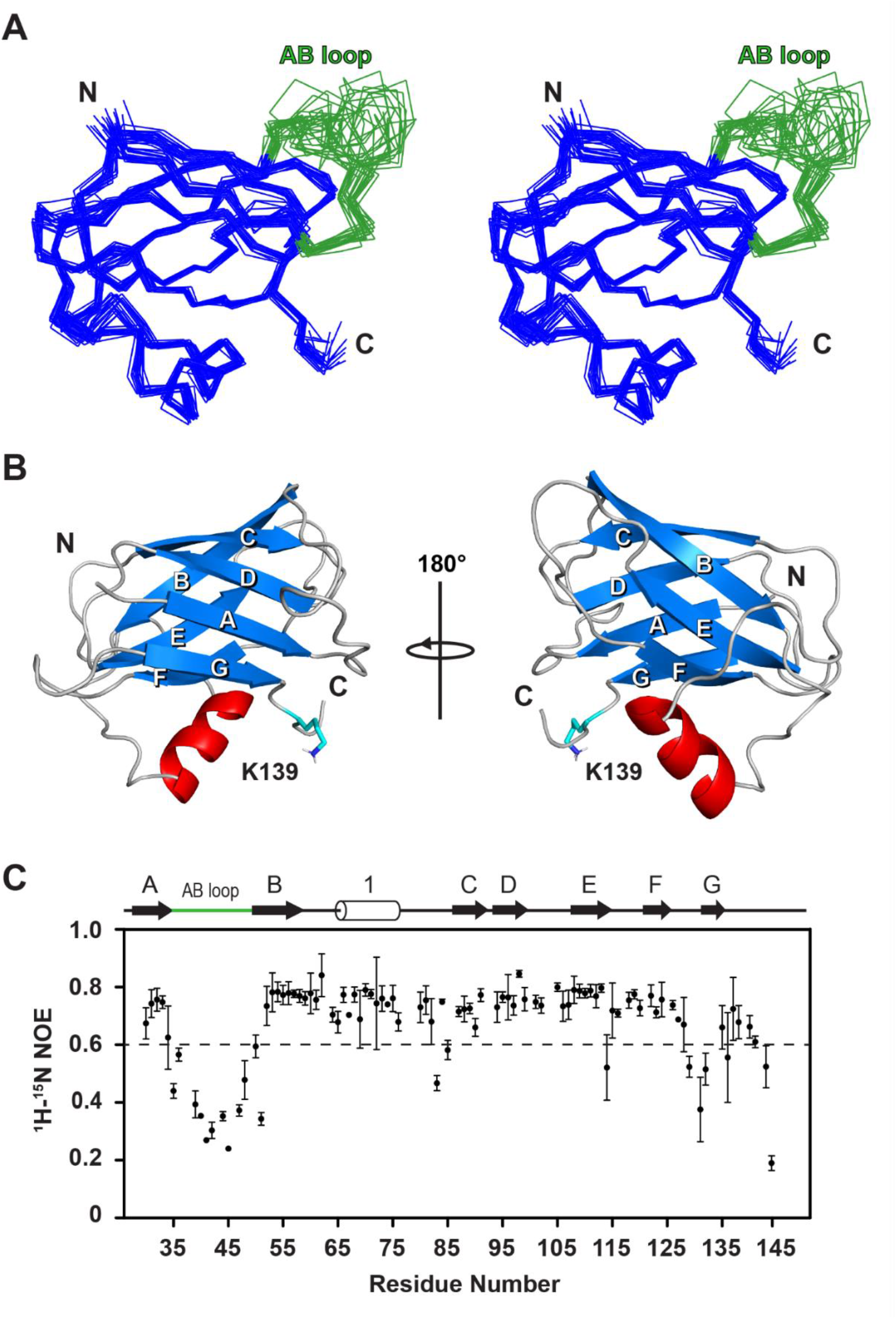
NMR structure of SpaB. (A) Stereoview showing the bundle of 20 lowest energy structures of SpaB. Residues that are structured in SpaB are colored blue, while residues in the disordered AB loop are colored green. Residues A27-D142 are shown. (B) Cartoon representation of the structure of SpaB with the strands of the β-sheet elements labeled A to G. The side chain of K139 that is crosslinked by the sortase enzyme is also shown. (C) Plot showing {^1^H}^15^N heteronuclear NOE (hetNOE) data for the backbone amide residues in SpaB. Flexible residues are located below the dashed line and have hetNOE values <0.6. The secondary structure is shown above the figure. The data was collected in triplicate and the error bars are the standard deviation of these measurements. The hetNOE data indicate that the AB loop is dynamic on the pico- to nano-second time scales.

An analysis of the structure using the program DALI identified 21 proteins in the Protein Data Bank with structural homology to SpaB (Z-scores > 4.0, **Table S3**). These proteins share only 6-21% sequence identity with SpaB, but nevertheless adopt related CnaB-type folds. 10 of the 21 structural homologs are pilin proteins that are expected to be crosslinked by pilus-specific sortase enzymes. The remainder of the homologs either have no known function or functions that are not related to pilus assembly (e.g., periplasmic copper-binding) and are not considered further. Several N-terminal domains within major pilins that are crosslinked by sortases are structurally related to SpaB, including FimP from *Actinomyces oris* (PDB: 3UXF) and the T6 backbone pilin from *S. pyogenes* (PDB: 4P0D). In addition to the SpaB protein from *C. diphtheriae*, structures of 5 basal pilins have been determined: RrgC (PDB: 4OQ1) from *S. pneumoniae*, FctB (PDB: 3KLQ) from *S. pyogenes*, GBS52 (PDB:3PHS) from *S. agalactiae*, and the SpaB (PDB:7CBS) and SpaE (PDB: 6JCH) basal pilins from *Lactobacillus rhamnosus* GG^24–28^ Interestingly, of these, only the basal pilins from *L. rhamnosus* share significant structural homology with SpaB. Moreover, only the basic fold is preserved among these structural homologs, and when they are compared to SpaB large differences are observed (the backbone atoms of the 21 homologs can only be aligned to SpaB with rmsd values ranging from 2.9-4.5 Å). Thus, our results are consistent with prior findings that have shown that pilins in gram-positive bacteria exhibit a high degree of structural variability^37^. Notably, SpaB is structurally related to ^N^SpaA which is also used by ^Cd^SrtA as a substrate^13^. A detailed comparison of their structures is presented in the Discussion section.

## Discussion

Covalently crosslinked pili in gram-positive bacteria mediate polymicrobial, host-pathogen, and host-commensal interactions^1-5^. These pili are produced via a biphasic process in which pilus-specific sortase enzymes first elongate the pilus by crosslinking its pilin components together via lysine-isopeptide bonds, followed by covalent attachment of the assembled pilus to the cell wall. The transition between pilin polymerization and cell wall anchoring is a key molecular switch that controls the length of the pilus, which has recently been shown to affect polymicrobial interactions^8,14^. Many pathogenic species of bacteria produce trimeric pili structures that contain specialized basal pilins that terminate pilus assembly after they are added to the base of the pilus (e.g. pili in *S. pyogenes, S. pneumoniae, S. agalactiae, E. faecalis*, and *C. diphtheriae* among others). In this study we investigated how the pilus-specific ^Cd^SrtA sortase terminates the assembly of the archetypal SpaA pilus from *C. diphtheria*e. We reconstituted the SpaB crosslinking reaction that adds this pilin to the base of the SpaA pilus to terminate its assembly and demonstrated that the ^Cd^SrtA enzymes connects SpaB to SpaA via a K139(SpaB)-T494(SpaA) isopeptide bond (**Fig. 2D**). This result is consistent with previous studies that demonstrated that K139 in SpaB is required for pilus assembly, and explicitly defines for the first time the chemical linkage that joins SpaB to the base of the pilus^38^. Furthermore, we show that amongst the three lysines in SpaB, only K139 is used as substrate by ^Cd^SrtA for crosslinking (**Fig. 2B,C**).

Measurements of pilin crosslinking and NMR experiments reveal that SpaB terminates pilus biogenesis by outcompeting SpaA for access to a shared ^Cd^SrtA-SpaA reaction intermediate (Scheme 1). On the cell surface the SpaA and SpaB pilins are thought to compete for access to ^Cd^SrtA. The shaft of the pilus is lengthened when the enzyme recognizes and crosslinks SpaA to the pilus, while pilus assembly is terminated when the enzyme crosslinks SpaB^12,14^. In these related crosslinking reactions, the ^Cd^SrtA enzymes recognizes lysine residues located within either the N-terminal domain of SpaA (^N^SpaA) or SpaB, which attack a shared ^Cd^SrtA-SpaA thioester intermediate in which ^Cd^SrtA’s active site C222 is joined to the C-terminal end of a second SpaA pilin (ES’ in scheme 1). Estimates of the steady-state kinetic parameters using a reconstituted system in which the pilins are crosslinked to a sorting signal peptide suggest that ^Cd^SrtA catalyzes the SpaB crosslinking reaction ∼6-times slower than ^N^SpaA crosslinking. Moreover, based on their K_M_ values, SpaB has ∼5-fold higher affinity for the ^Cd^SrtA-SpaA intermediate as compared to ^N^SpaA (^SpaB^K_M_ and ^SpaA^K_M_ values of 3.2 ± 0.2 µM and 16 ± 4 µM, respectively). Additional evidence that supports the idea that the pilins compete for access to the ^Cd^SrtA-SpaA thioester intermediate comes from competition experiments in which an unreactive SpaB^K139A^ pilin was added to the ^N^SpaA crosslinking reaction (**Fig. 4C**). SpaB^K139A^ inhibits ^N^SpaA crosslinking in a manner consistent with its binding to the ^Cd^SrtA-SpaA thioester intermediate with a K_D_ ∼65 µM. NMR experiments did not detect significant interactions between SpaB and ^Cd^SrtA (**Fig. 5**). Thus, we conclude that SpaB only binds ^Cd^SrtA after it has reacted with SpaA to create the ^Cd^SrtA-SpaA intermediate and the SpaA and SpaB pilins compete for access for this enzyme-substrate complex. On the cell surface this competition may also involve the membrane embedded SrtA and SrtF enzymes, which form thioester enzyme-substrate complexes with SpaA and SpaB, respectively.

The NMR structure and dynamics of SpaB are strikingly similar to ^N^SpaA, suggesting that ^Cd^SrtA recognizes similar features within these pilin substrates (**Fig. 7B**). Despite sharing only 16% primary sequence identity, both proteins adopt a CnaB-type fold and possess similar secondary structural topologies (**Fig. 7A**). In each pilin the reactive lysine is located within a polypeptide segment that immediately follows the last β-strand (strand G) (K190 in ^N^SpaA and K139 in SpaB). SpaB and ^N^SpaA also share another landmark feature of many gram-positive pilin proteins, the presence of an extended disordered loop that connects β-strands A and B (AB loop), which is positioned adjacent to their reactive lysine residues (colored green in **Fig. 7C**). The AB loops in SpaB and ^N^SpaA are flexible on the pico- to nano-second time scales based on {^1^H}^15^N heteronuclear NOE data (**Fig. 6C**), but the loop in SpaB is positioned farther away from the reactive lysine as compared to ^N^SpaA^10^. The backbone coordinates of SpaB and ^N^SpaA can only be superimposed with a RMSD of 3.8 Å (**Fig. 7B**). Visual inspection reveals that although the β-sheet conformations are largely preserved between the two modules, significant differences occur in the conformations and lengths of the loops that connect the strands of the β-sheet, including the presence of an additional α-helix connecting strands B to C in ^N^SpaA (**Fig. 7A**).

**Figure 7.**
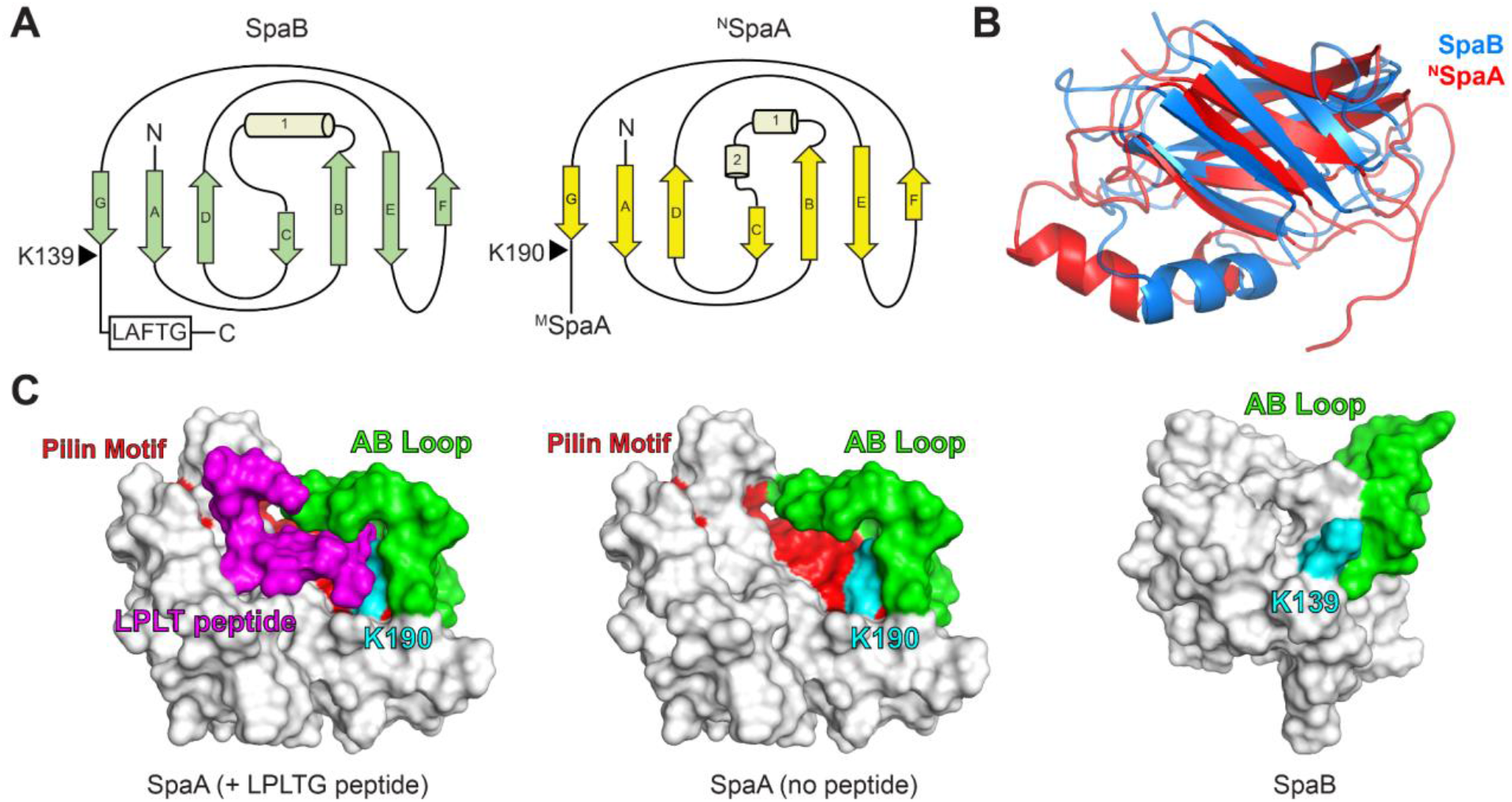
Comparison of SpaB and ^N^SpaA. (A) Schematic representation of the secondary structure topologies of SpaB and ^N^SpaA. The β-strands are designated A through G and the helices are numbered. The reactive lysine and sorting signals are shown. (B) Overlay of the backbone structures of SpaB (blue) and the ^N^SpaA (red) (PDB: 3HR6). (C) Comparison of the structures of ^N^SpaA and SpaB. The surface representation of ^N^SpaA crosslinked to its sorting signal peptide is shown on the left (PDB: 7K7F). The structures of the apo-forms of ^N^SpaA and SpaB are shown in the middle and right panels, respectively. Color code: AB loop (green), peptide (magenta), reactive lysine (cyan) and pilin motif residues (red).

The SpaA-x-SpaB and SpaA-x-SpaA crosslinked interfaces can be expected to be structurally distinct based on the NMR structure of SpaB and previous studies of the crosslinked SpaA-x-SpaA interface. The structure of ^N^SpaA crosslinked to a KNAGFELPLT sorting signal peptide derived from the C-terminal end of SpaA (^N^SpaA-LPLT complex) has been determined^10^. This work revealed that lysine-isopeptide bond formation triggers a disordered-to-ordered transition in ^N^SpaA’s AB loop that partially masks the K190(^N^SpaA)-T494(^C^SpaA) linkage to stabilize the SpaA-x-SpaA interface (**Fig. 7C**, left). In the complex, amino acids AGFELPLT within the bound peptide rest in a groove that is positioned at the edge of the β-sandwich. The floor of the binding groove is formed by residues in strands F and G. The peptide in the complex is positioned in a nearly antiparallel manner with respect to strand F, enabling its N-terminus to exit near the BC and FG loops that are predicted to form an interpilin interface within the pilus fiber. This binding groove contains residues in the conserved WxxxVxVYPK pilin motif that are located in strand G (**Fig. 7C**, middle colored red). Even though its primary sequence does not contain a pilin motif, the structure of SpaB does possess a similar binding groove for the crosslinked sorting signal, but it is much smaller and can presumably only accommodate the side chains of the Leu-Pro-Leu residues in the sorting signal. Signal binding to this site presumably requires that the peptide bind in a distinct orientation relative to what is seen in the ^N^SpaA-LPLT complex in order to accommodate the closer proximity of SpaB’s helix to the reactive K139 residue. Also notable is the lack of primary sequence conservation between SpaB and ^N^SpaA in their BC and FG loops that are predicted to form the interface between ^C^SpaA in both the SpaB-x-SpaA and SpaA-x-SpaA interfaces. Deeper insight into the structure of the SpaA pilus and its assembly mechanism will require resolving the atomic structure of these interfaces and learning how SpaB and ^N^SpaA are recognized by ^Cd^SrtA to initiate crosslinking. As gram-positive pili are key virulence factors, this knowledge could lead to the development of novel antibiotics that work by disrupting pilus biogenesis.

## Acknowledgements

This paper is dedicated to G. Marius Clore. We thank him for his inspiring mentorship and support, as well as his many contributions to the field of structural biology. We also thank members of the Clubb group for useful discussions. This research was supported by grants from the National Institutes of Health (NIH) (AI52217 [to R. T. C.]; GM103479, GM145286 [to J. A. L.]) T32GM007185 [to C. K. S.]; T32GM145388 [to N. A. C.]. J. Y. F. acknowledges the support from the National Science Foundation (NSF) Graduate Research Fellowship program (DGE-1650604).

