## Supplemental Information for "The Basal and Major Pilins in the *Corynebacterium diphtheriae* SpaA Pilus Adopt Similar Structures that Competitively React with the Pilin Polymerase"

Table S1: Masses identified in the SpaB crosslinked complex

Table S2: Statistics table for the NMR structure of SpaB

Table S3: DALI analysis for the structure of SpaB

**Table S1. Tandem mass spectrometry analysis of SpaA-SpaB isopeptide linkage**

| Fragment Ion | Charge | Observed *m/z* | Theoretical *m/z* | Difference between observed and theoretical (ppm) |
|---|---|---|---|---|
| Phenylalanine Immonium | 1+ | 120.0805 | 120.0813 | -6.995 |
| Lysine Marker Ion | 1+ | 129.1021 | 129.1022 | -1.0070 |
| $a°_{2\alpha}$ | 1+ | 181.1328 | 181.1335 | -3.9750 |
| $b°_{2\alpha}$ | 1+ | 209.1273 | 209.1285 | -5.6425 |
| $a_{2\beta}$ | 1+ | 249.1231 | 249.12341 | -1.3246 |
| $b_{2\beta}$ | 1+ | 277.1177 | 277.11833 | -2.2373 |
| $b_{3\beta}$ | 1+ | 390.2027 | 390.2023 | 0.9482 |
| $M_{\alpha}/y_{3\beta}$ | 2+ | 391.2604 | 391.2627 | -5.8528 |
| $b_{2\alpha}$ / $y_{1\beta}$ | 1+ | 440.2829 | 440.2867 | -8.5626 |
| $M_{\alpha}$ | 1+ | 470.3286 | 470.3337 | -10.8859 |
| $b_{2\alpha}$ / $y_{3\beta}$ | 1+ | 537.3351 | 537.3395 | -8.2257 |
| $M_{\alpha}/y_{1\beta}$ | 1+ | 571.3779 | 571.3814 | -6.1780 |
| $b_{2\alpha}$ / $y_{4\beta}$ | 1+ | 650.42 | 650.4236 | -5.5656 |
| $M_{\alpha}/y_{2\beta}$ | 1+ | 684.4686 | 684.4654 | 4.71901 |
| $M_{\alpha}/y_{3\beta}$ | 1+ | 781.5147 | 781.5182 | -4.4657 |
| $M_{\alpha}/y_{4\beta}$ | 1+ | 894.5938 | 894.6023 | -9.4902 |

**Table S2. Structural statistics of the solution structure of SpaB**

| | ⟨SA⟩[a] |
|---|---|
| Root mean square deviation | |
| NOE Interproton distance restraints (Å) | |
| All (1277) | 0.036 ± 0.003 |
| Dihedral angles restraints (°)[c] (192) | 0.73 ± 0.08 |
| Deviation from idealized covalent geometry | |
| bonds (Å) | 0.0110 ± 0.00004 |
| angles (°) | 0.73 ± 0.01 |
| impropers (°) | 0.036 ± 0.02 |
| PROCHECK results (%) | |
| most favorable region | 71.0 ± 2.4 |
| additionally allowed region | 24.9 ± 3.0 |
| generously allowed region | 4.1 ± 1.3 |
| disallowed region | 0.0 ± 0.0 |
| Coordinate Precision (Å) | |
| Protein backbone | 0.56 ± 0.06 |
| Protein heavy atoms | 1.51 ± 0.35 |

**Table 3. DALI Analysis of SpaB**

| Sl. No. | Chain | Z-score | RMSD | Alignment Length (aa) | No. of Residues (aa) | Sequence Identity (%) | PDB | Description |
|---|---|---|---|---|---|---|---|---|
| 1 | A | 6.4 | 4.1 | 101 | 449 | 17 | 3uxf | Structure of the fimbrial protein FimP from *Actonomyces oris* |
| 2 | A | 6.3 | 3.2 | 99 | 470 | 17 | 4p0d | Structure of the T6 backbone pilin of serotype M6 *Streptococcus pyogenes* |
| 3 | A | 5.8 | 3.8 | 92 | 143 | 14 | 7k7f | Structure of the *Corynebacterium diphtheriae* SpaA Pilin-Signal Peptide Complex |
| 4 | A | 5.8 | 3.6 | 93 | 132 | 17 | 7cbs | Structure of SpaB basal pilin from *Lactobacillus rhamnosus* GG |
| 5 | A | 5.1 | 3.3 | 70 | 522 | 21 | 6fwv | Structure of the *Bacillus anthracis* TIE protein |
| 6 | A | 5 | 3.2 | 92 | 331 | 11 | 4oq1 | Structure of the *Ancillary Pilin RrgC of Streptococcus pneumoniae* |
| 7 | E | 5 | 3.4 | 73 | 202 | 11 | 4fxt | Structure of a DUF3823 family protein from *Bacteroides ovatus* ATCC |
| 8 | A | 4.8 | 3.6 | 75 | 242 | 16 | 4eiu | Structure of a DUF3823 family protein (BACUNI_03093) from Bacteroides uniformis ATCC 8492 |
| 9 | B | 4.5 | 3.4 | 80 | 161 | 13 | 4gqz | Structure of the periplasmic copper-binding protein CueP from S*almonella enterica* serovar Typhimurium |
| 10 | A | 4.5 | 4.4 | 96 | 327 | 14 | 6jch | Structure of SpaE basal pilin from *Lactobacillus rhamnosus* GG |
